# genomeSidekick: a user-friendly epigenomics data analysis tool

**DOI:** 10.1101/2022.04.18.488677

**Authors:** Junjie Chen, Ashley J. Zhu, René R. Sevag Packard, Thomas M. Vondriska, Douglas J. Chapski

## Abstract

Recent advances in epigenomics measurements have resulted in a preponderance of genomic sequencing datasets that require focused analyses to discover mechanisms governing biological processes. In addition, multiple epigenomics experiments are typically performed within the same study, thereby increasing the complexity and difficulty of making meaningful inferences from large datasets. One gap in the sequencing data analysis pipeline is the availability of tools to efficiently browse genomic data for scientists that do not have bioinformatics training. To bridge this gap, we developed genomeSidekick, a graphical user interface written in R that allows researchers to perform bespoke analyses on their transcriptomic and chromatin accessibility or chromatin immunoprecipitation data without the need for command line tools. Importantly, genomeSidekick outputs lists of up- and downregulated genes or chromatin features with differential accessibility or occupancy; visualizes ‘omics data using interactive volcano plots; performs Gene Ontology analyses locally; and queries PubMed for selected gene candidates for further evaluation. Outputs can be saved using the user interface and the code underlying genomeSidekick can be edited for custom analyses. In summary, genomeSidekick brings wet lab scientists and bioinformaticians into a shared fluency with the end goal of driving mechanistic discovery.

## Introduction

Computational biology tools written in different languages and applied across diverse fields allow for creative interrogation of genomics data to make biological conclusions. Understandably, the breadth of online genomic data analysis resources may appear overwhelming to a novice programmer. Fortunately, global efforts to bring bioinformatics training to general researchers are well underway (Mulder et al., 2018). Nevertheless, learning how to code may be a barrier to entry for non-bioinformaticians into the field of epigenomics, yet it is important to incorporate these researchers into the data analysis process. A logical solution to this training issue is an inclusive approach that brings non-bioinformaticians into computational workflows after completing most of the command line processes, thereby fostering scientific creativity and leveraging shared knowledge about how the data are processed, analyzed, and visualized.

While lab skillsets ideally include formal bioinformatics knowledge, genomic researchers who do not understand how to code can readily make meaningful conclusions using processed data. An unmet need within this realm is a software for visualizing genomics data and filtering epigenomic and transcriptomic results for downstream analyses, especially considering the combination of orthogonal genomic datasets required to reveal more comprehensive mechanisms of cell biology. In addition, while Excel is a common tool for management and visualization of data, gene lists can be imported incorrectly into Excel and cause permanent edits to gene names (Ziemann et al., 2016). To prevent this issue and to promote independence from the bioinformatician, the next logical step is to furnish tools to perform data operations that a novice researcher might otherwise try in Excel.

The availability of distinct measurements to understand genomic mechanisms governing complex cellular and organ phenotypes has increased over time, resulting in a need to combine datasets (Chapski and Vondriska, 2021). Our recent study using RNA-seq, ATAC-seq, reduced representation bisulfite sequencing (RRBS), and chromatin structural data is an example of such integration of orthogonal data to make meaningful conclusions about chromatin architectural dynamics during heart failure (Chapski et al., 2021). Another investigation established an atlas of murine ATAC-seq and RNA-seq data across 86 immune cell types and integrated the two datasets to identify a subset of cell types containing open regulatory elements bound by retinoic acid receptor-related orphan receptor gamma (RORγ) or paired-box protein PAX5 (as measured by ChIP-seq), thereby linking chromatin accessibility, transcription, and transcription factor binding in specific cell types (Yoshida et al., 2019). A common feature of all ‘omics investigations is the need to ask questions of the massive datasets once acquired—to prioritize for further mechanistic evaluation. We also appreciate that even professional bioinformaticians may not have the time to perform bespoke analyses for collaborators: thus, a tool for transforming lists of genes into functional targets for a focused, mechanistic experiment is an opportunity to bring non-computational scientists and clinicians into the genomic analysis process.

To bridge the gap between processed data and biological inference, we built user-friendly genomic data visualization and manipulation tools for investigators without computational training. This GUI-based software called genomeSidekick allows for investigation of transcriptomic (RNA-seq) data in addition to chromatin accessibility (ATAC-seq) and chromatin immunoprecipitation-sequencing (ChIP-seq) data in a web browser. Based on a Shiny (Chang et al., 2021) dashboard written in R (Team, 2020), our tool—which we have named genomeSidekick—generates commonly used, intuitive graphs with interactive information retrieval. Moreover, we wrapped data visualization features for each individual experiment (RNA-seq, ATAC-seq, and ChIP-seq) into individual tabs to make switching between experiments easier. We also provide a tab to integrate RNA-seq, ATAC-seq and ChIP-seq datasets, so modulation of the transcriptome and epigenome can be examined based on multiple criteria from independent experiments. Lastly, we provide links to external tools (and offer to perform small analyses locally) to facilitate Gene Ontology analysis and PubMed searches.

Freely available on GitHub (https://www.github.com/dchapski/genomeSidekick), genomeSidekick also contains extensive user-friendly documentation in a README markdown file with informational links so that most novice bioinformaticians can achieve results quickly. Lastly, genomeSidekick is a customizable tool that allows for code editing to support a shared collaboration between bioinformaticians and non-computational personnel in the biological research setting, thereby promoting increased computational engagement by non-bioinformaticians.

## Methods

To run genomeSidekick, users should download the software from the repository on GitHub (https://www.github.com/dchapski/genomeSidekick) and then open the app.R file using Rstudio and click the “Run” button in the upper right corner of the script. Alternatively, users can download the code and run the app directly from the terminal using “R -e “shiny::runApp(“/path/to/app.R”)”. Comma-separated or tab-delimited input RNA-seq data should include gene names (either identifiers or common names) with an adjusted p-value and log2FoldChange (preferably from a tool such as DESeq2 (Love et al., 2014) or edgeR (Robinson et al., 2010), which correct p-values for multiple testing and provide fold change information). Comma-separated or tab-delimited input ATAC-seq data should include adjusted p-values and log2FoldChange information about accessibility peaks (the output from DiffBind (Ross-Innes et al., 2012) works well), in addition to either the closest gene or an overlapping gene for each feature. Gene names should be included for the ATAC-seq data as they are required for merging the RNA-seq and ATAC-seq dataset; however, independent analysis of ATAC-seq data alone does not require gene names. Importantly, other epigenomic experiments such as ChIP-seq outputs containing log2FoldChanges and adjusted p-values can be used on the genomeSidekick platform, either alone or in combination with RNA-seq data as described above for the ATAC-seq tab.

Extensive documentation regarding installation of R, RStudio, and dependencies for genomeSidekick is provided on the GitHub page. This documentation is also provided within the software so users can directly find information on how to run the software within the app. To run the app in a password protected location online, a Shiny subscription can be purchased from the RStudio website (pricing starts at $9 USD/month in 2021). For exploration, we also provide online access at https://genomesidekick.shinyapps.io/genomesidekick/.

## Results

The genomeSidekick software, written in R, can be run on a laptop and requires few dependencies to analyze RNA-seq, ATAC-seq, ChIP-seq, and any other epigenomics datasets that contain p-values and fold changes. The utility of genomeSidekick comes from its easy-to-use interface built on the Shiny framework in R (Team, 2020). Shiny is an R package that provides a front- and back-ends for user-friendly investigation of datasets. In addition, Shiny apps can be hosted online to facilitate data sharing and exploration. This genomics dashboard allows separation of experimental strategies via individual tabs in the GUI (**Figure 1**). Inputs include processed data tables that can be loaded directly into the app. For example, an output from DESeq2 (Love et al., 2014) that contains the log2FoldChange and adjusted p-value information required for the visualizations. Other inputs include the output from DiffBind (Ross-Innes et al., 2012), a tool that statistically evaluates differentially bound or accessible genomic regions in the case of chromatin immunoprecipitation followed by sequencing (ChIP-seq) or ATAC-seq data, respectively.

**Figure 1.**
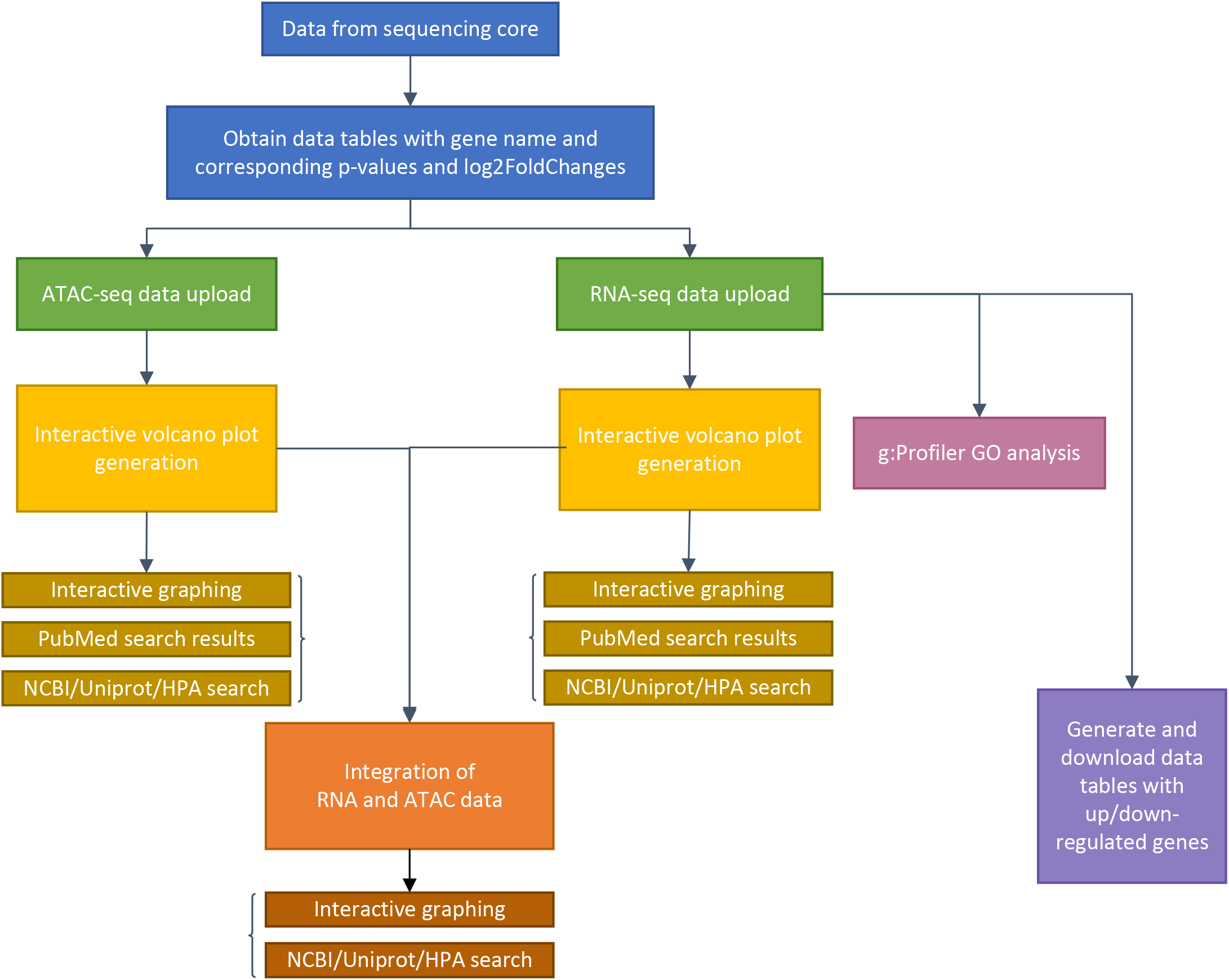
Flow chart detailing genomeSidekick functionalities. After raw sequencing data is obtained from the sequencing core, alignment and statistical analyses on genes or chromatin features should be performed by a bioinformatician, with at least three columns as a result: feature name, adjusted p-values, and log2FoldChange (*blue boxes*). Then, genomeSidekick users can upload these data tables (*.csv or tab-delimited files*) as inputs to their corresponding tabs within the Shiny app (*green boxes*), which will contain functions for generation of independent interactive volcano plots (*yellow boxes*) as well as integrated ATAC-seq and RNA-seq graphs (*orange boxes*), for example. Note that in addition to ATAC-seq, users may analyze ChIP-seq and other epigenomics data inputs containing p-values and log2FoldChanges. Uploading an RNA-seq data set will also allow the application to perform g: Profiler GO analysis (*magenta box*) and generate downloadable lists of of up- and downregulated genes (purple box). HPA: Human Protein Atlas.

Visualizations for volcano plots are coded using ggplot2 (Wickham, 2016) based code and visualized using ggplotly (Sievert, 2020), an open-source R package allows for interactive inspection of graphs (**Figure 2A-C**). The ggplotly visualizations allow for truly interactive point-by-point investigation to reveal individual metrics about each data point (gene name, adjusted p-value, log2FoldChange, and other custom information within the table). Superimposed on these visualizations are gene names highlighted by small lines (visualized using ggrepel (Slowikowski, 2021)) to indicate the *n* most significant points in the dataset. Notably, when a gene point is clicked within a volcano plot, genomeSidekick links the user to either the NCBI database, the UniProt (UniProt, 2021) website, or the Human Protein Atlas (Uhlen et al., 2015) for further investigation of candidate gene functions. Some genes are not available for data visualization since many tools that calculate differential expression/accessibility only statistically evaluate loci containing experimental data, thereby resulting in unmeasured regions without a p-value (**Figure 2D** shows an example of this phenomenon). When RNA-seq and ATAC-seq (or ChIP-seq) inputs include common genomic feature information (for example, gene names), genomeSidekick can merge and filter these tables to produce a list of the *n* most upregulated and downregulated genes with accessibility information. In addition, the merge computation is performed in a way that gene names do not become corrupted from loading data in Excel (for more, see Introduction above and (Ziemann et al., 2016)). This merged dataset can then be visualized as a volcano plot: one dataset (RNA-seq, for example) is plotted along the axes and the other dataset visualized using different point size and coloring to show addition information (**Figure 2E**).

**Figure 2.**
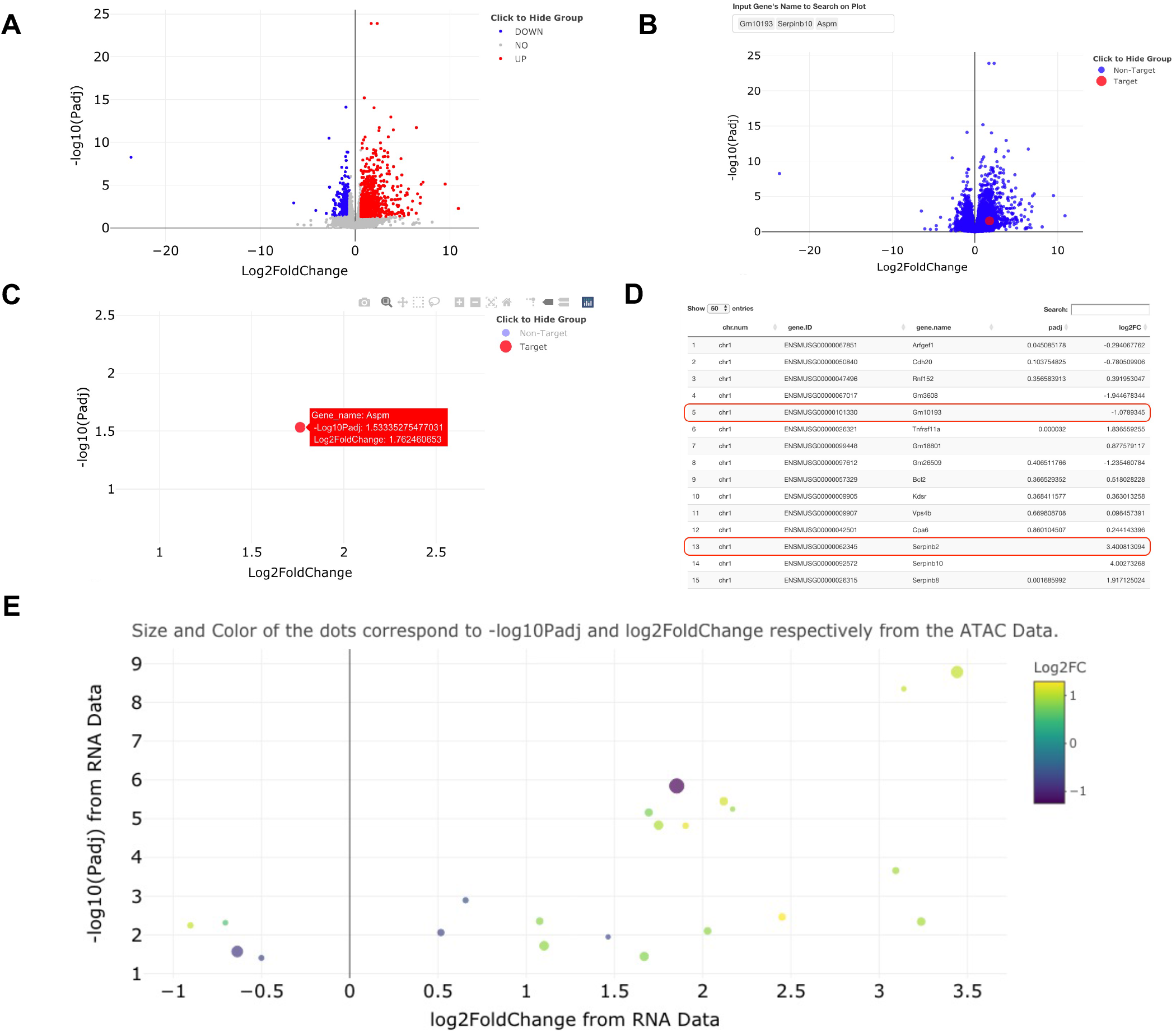
Example visualizations of sample RNA-seq and ATAC-seq data using genomeSidekick. **(A)** Volcano plot of RNA-seq data shows down-regulated genes in blue, up-regulated genes in red, and nonsignificant genes in gray. Thresholds for p-value and log2FoldChange can be adjusted within genomeSidekick. **(B)** Searching for a particular gene(s) will highlight the target gene in red and mark the non-target genes in blue. In the example query for three genes, only one is found and visualized on the graph. **(C)** The non-target genes can be hidden by clicking on the Non-Target label in the legend (resulting in a grey label) and un-hidden by clicking the label a second time. Details about any gene shown on the interactive volcano plot can be shown by hovering over the dot on the plot. **(D)** Looking at the RNA-seq data upload page, genes without a p-value are not plotted on the volcano plot and therefore will not return any results when queried on the graph (see red lines). Common RNA-seq analysis packages do not evaluate all genes in the genome due to low detection, and this varies by experiment. Panels A through D use RNA-seq data as an example, but the functionalities are the exact same for ATAC-seq data. **(E)** Once both RNA-seq and ATAC-seq data sets are uploaded, they can be integrated into one graph using either the RNA-seq or ATAC-seq data as the base. Example data shown have padj < 0.05 and FDR < 0.1 in the RNA and ATAC-seq datasets, respectively.

To test the ease of dataset integration in a setting outside our institution, a collaborator provided a use case for custom analysis of RNA-seq and ATAC-seq data from (Chapski et al., 2021). Specifically, this collaborator sought to determine how the expression and chromatin accessibility at gene loci change with 3 days cardiac pressure overload (a pathological model that eventually leads to heart failure) in mice. Interestingly, the integrated RNA-seq and ATAC-seq output of genomeSidekick showed a significant increase in transcription and chromatin accessibility at the *Xirp2* gene locus (**Figure 3**), consistent with an earlier study showing that the cardiac stressor angiotensin II elicits an increase in *Xirp2* transcription mediated by the transcription factor MEF2A (McCalmon et al., 2010). This exercise, performed on the collaborator’s first exploration of the software, suggests that genomeSidekick is useful for quick exploration of datasets and can provide meaningful scientific insights to first time users.

**Figure 3.**
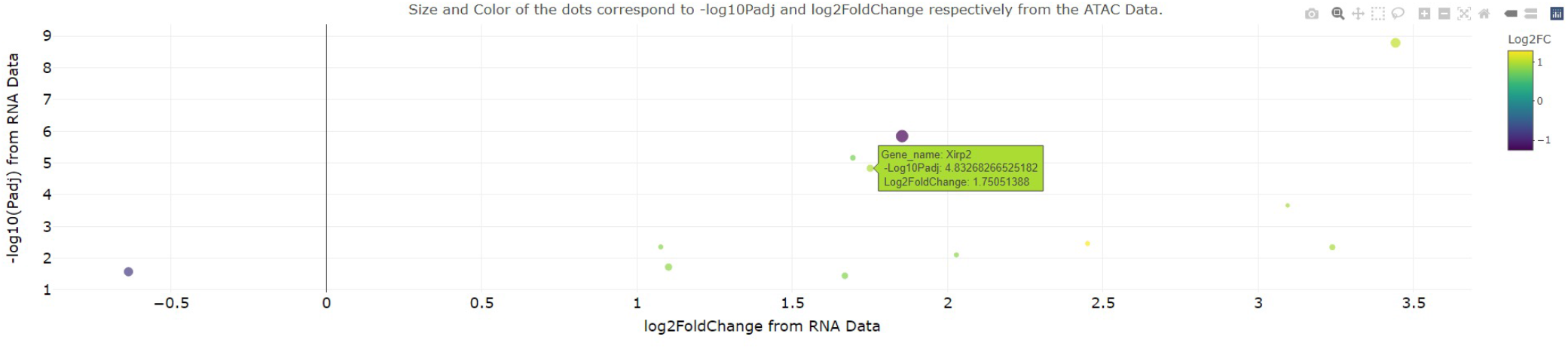
Example integrated RNA-seq/ATAC-seq visualization from a collaborator’s first time using genomeSidekick. A collaborator recently visualized change in transcription and promoter accessibility at the murine *Xirp2* locus after 3 days cardiac pressure overload. This gene, also upregulated at the transcriptional level after angiotensin II treatment, also becomes more accessible with pressure overload, suggesting that the observed increase in transcription is a consequence of increased accessibility. Dot size and color scale indicate ATAC-seq –log10(adjusted p-value) and log2FoldChange, respectively. Example data shown have padj and FDR less than 0.05 in the RNA and ATAC-seq datasets, respectively.

Importantly, the gene list outputs from each genomeSidekick tab are displayed for direct use as inputs for other software. For example, genomeSidekick includes a feature to perform local Gene Ontology analysis on smaller gene list outputs using the gprofiler2 (Kolberg et al., 2020) package in R in addition to a link to the g:Profiler website (Raudvere et al., 2019) for analyses of larger output gene lists from genomeSidekick that might take longer on a local machine. Lastly, we include a feature for quick PubMed searches of genes of interest that outputs query results directly in the app. This feature is based on the easyPubMed package in R (Fantini, 2019) and is designed to keep users’ eyes on their data instead of opening a new tab to perform queries on data points of interest. Taken together, these features allow a non-bioinformatician to increase their computational fluency without having to learn how to code.

## Discussion

We built a tool called genomeSidekick to facilitate inclusion of non-bioinformaticians into computational workflows for RNA-seq, ATAC-seq, ChIP-seq, or any other datasets that undergo statistical testing. This GUI-based software written in R allows individuals to focus their efforts on biological inference without having to frontload the bioinformatics training required to maneuver the command line. Specifically, genomeSidekick facilitates integration of gene expression and chromatin accessibility data, for example, to narrow down gene lists for further analyses. In addition, the software provides an opportunity for non-bioinformaticians to perform small edits to the code to customize their visualizations and filtering criteria without having to learn R. Overall, genomeSidekick will bring wet lab scientists onto a more level playing field for common data analysis questions, thereby reducing dependence on bioinformaticians.

Additional software exists for analysis of gene expression and epigenomics data and may be useful for more computationally versed individuals. For example, DEApp can be used to perform differential expression testing and data visualization (Li and Andrade, 2017), although a significant hurdle to using this tool is knowing which statistical approach to use for differential expression within the software. In addition, DEBrowser (Kucukural et al., 2019) and VisRseq (Younesy et al., 2015) are useful for performing end-to-end bioinformatics analyses of datasets, and both programs complete complicated tasks such as heatmap generation and principal component analysis. Importantly, these tools may require knowledge of data transformations at each step of a given analysis and/or training in statistics. In contrast, genomeSidekick provides a platform for users to explore and integrate processed transcriptomics and epigenomics data and create figures without the complexity seen in other tools.

The simplicity of genomeSidekick allows researchers with no bioinformatics background to investigate their own datasets after initial mapping, quantification, and differential testing by a bioinformatician. Thresholding of p-values for individual experiments can be edited for custom stringency, which allows wet lab researchers to perform independent analyses without requesting individual gene lists from a bioinformatician. Moreover, extensive documentation providing explanations of individual functions and links to learning resources is condensed into an intuitive README file on GitHub with an intuitive interface and examples.

The genomeSidekick application allows bioinformaticians to send data to collaborators and then have them interact with multiple datasets independently. Importantly, the app can be hosted online for a small monthly fee using https://www.shinyapps.io, thereby facilitating longer distance collaborations. Accordingly, for simple data exploration, we provide genomeSidekick online at https://genomesidekick.shinyapps.io/genomesidekick/. Overall, genomeSidekick will help bring wet lab researchers into the computational realm by fostering creativity with data visualization and integrative analyses in a user-friendly format.

## Acknowledgements

The authors would like to thank collaborator Dr. Christoph D. Rau for software testing and members of the Vondriska Lab for comments and suggestions. Research in the Vondriska Lab is supported by the NIH, UCLA Clinical and Translational Science Institute, the Department of Anesthesiology & Perioperative Medicine, and the David Geffen School of Medicine at UCLA. Dr. Packard is supported by VA Merit BX004558 and the UCLA Cardiovascular Discovery Fund/Lauren B. Leichtman and Arthur E. Levine Investigator Award.

## Author Contributions

JC and DJC conceived of the study. JC, AJZ, and DJC wrote the software. RRSP and TMV provided infrastructure. DJC supervised the project and wrote the manuscript. All authors read and approved the final manuscript.

